# Bimodal regulation of axonal transport by the GDNF-RET signalling axis in healthy and diseased motor neurons

**DOI:** 10.1101/2022.04.26.489550

**Authors:** Elena R Rhymes, Andrew P Tosolini, Alexander D Fellows, William Mahy, Neil McDonald, Giampietro Schiavo

## Abstract

Deficits in axonal transport are one of the earliest pathological outcomes in several models of amyotrophic lateral sclerosis (ALS), including SOD1^G93A^ mice. Evidence suggests that rescuing these deficits prevents disease progression, stops denervation, and extends survival. Kinase inhibitors have been previously identified as transport enhancers, and are being investigated as potential therapies for ALS. For example, inhibitors of p38 mitogen-activated protein kinase and insulin growth factor receptor 1 have been shown to rescue axonal transport deficits *in vivo* in symptomatic SOD1^G93A^ mice. In this work, we investigated the impact of RET, the tyrosine kinase receptor for glial cell-line-derived neurotrophic factor (GDNF), as a modifier of axonal transport. We identified fundamental interplay between RET signalling and axonal transport in both wild type and SOD1^G93A^ motor neurons *in vitro*. We demonstrated that blockade of RET signalling using pharmacological inhibitors and genetic knockdown enhances signalling endosome transport in wild type motor neurons and uncovered a divergence in the response of primary motor neurons to GDNF compared with cell lines. Finally, we demonstrated that inhibition of the GDNF-RET signalling axis rescues *in vivo* transport deficits in early symptomatic SOD1^G93A^ mice, promoting RET as a potential therapeutic target in the treatment of ALS.

## Introduction

Axonal transport, the process by which mRNAs, protein complexes and organelles move between synaptic terminals and the soma, is fundamental for neuronal survival ^1^. Deficits in this process occur in many neurodegenerative diseases, including amyotrophic lateral sclerosis (ALS)^2,3^. Anterograde transport towards axonal terminals is mediated by members of the kinesin motor protein superfamily, whereas retrograde transport is driven by cytoplasmic dynein. Human motor neuron (MN) axons can be over a meter long, necessitating efficient axonal transport. Accordingly, an impairment in axonal transport is one of the earliest pathological outcomes observed in several models of amyotrophic lateral sclerosis (ALS)^4–7^.

Signalling endosomes are generated following the activation and internalisation of neurotrophin receptors, such TrkB and p75^NTR^, and predominantly undergo retrograde axonal transport ^8^. Analysis of signalling endosome dynamics in the sciatic nerve of SOD1^G93A^ mice, an established animal model of ALS, revealed a significant slowing of retrograde transport compared to wild type (WT), both at pre-symptomatic and symptomatic disease stages^7,9^. Axonal transport deficits have also been observed in primary MN cultures from SOD1^G93A^ (Ref 5) and TDP-43^10^ embryos, cultured SOD1^G93A^ spinal cord explants^11^, induced pluripotent stem cells (iPSCs)-derived MNs harbouring ALS causative mutations in SOD1^12^, FUS^13^ and TDP-43 (Ref 14), as well as *in vivo* in transgenic TDP-43^M337V^ mice^6^. Therefore, rescuing these early deficits is a key priority for therapeutics in ALS.

Protein kinases regulate axonal transport^15^ and as such, kinase inhibitors represent a class of pharmacological compounds with the potential to boost axonal transport. While until recently the main focus of translational efforts has been the role of kinases in cancer^16^, kinase overactivation has also been identified in neurological diseases^17^, and linked to alterations in axonal transport. For example, GSK3-β has been shown to phosphorylate kinesin and induce a significant slowing of anterograde axonal transport^18^. Interestingly, aberrant kinesin phosphorylation occurs after treatment with mutant SOD1 protein^19^. GSK3-β has also been shown to inhibit retrograde transport through phosphorylation of the dynein intermediate chain, reducing dynein’s force generation^20^. Many kinase inhibitors have been FDA-approved for cancer treatment^16^, offering the possibility for repurposing already-approved drugs for neurodegenerative diseases, offering significantly faster patient-access time. Importantly, rescuing axonal transport deficits can significantly improve ALS phenotypes. For example, inhibitors of histone deacetylase 6 (HDAC6) have been shown to increase axonal transport speeds^13^, and HDAC6 deletion in SOD1^G93A^ mice increases neuromuscular junction innervation and extends survival^21^.

In line with this, we performed a pilot screen by treating mouse embryonic stem cell-derived (ES) MN with a small kinase inhibitor library, and identified several potential enhancers of axonal transport (4, results shown in Fig. 1A). Two of these hits have since been validated, with acute inhibition of both p38 mitogen-activated protein kinase (MAPK) and insulin-like growth factor 1 receptor (IGF1R) rescuing retrograde axonal transport deficits in SOD1^G93A^ MN both *in vitro* and *in vivo*^4,22^.

**Figure 1.**
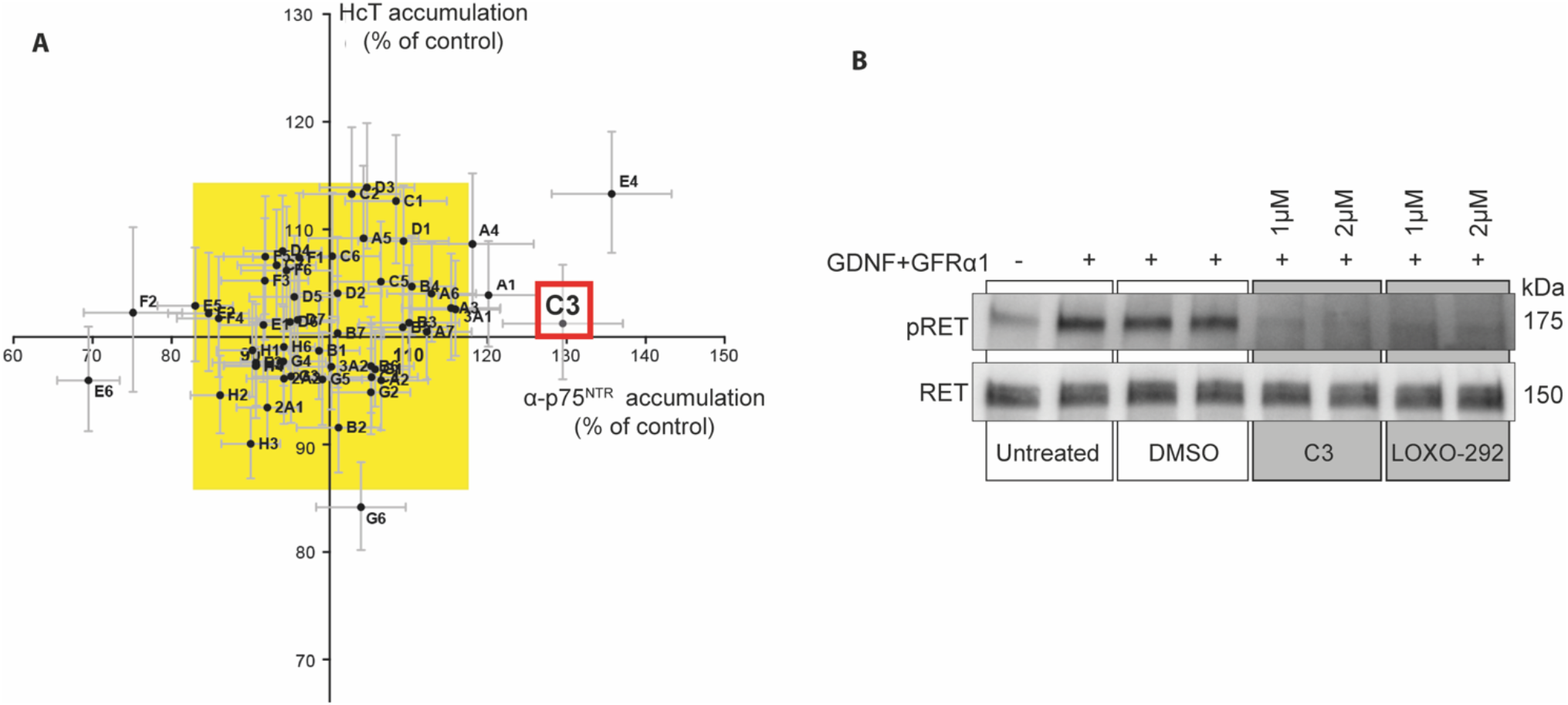
A kinase inhibitor screen identified C3, a RET inhibitor, as an enhancer of axonal transport. **A**. Summary of results from a kinase inhibitor pilot screen ^4^. The graph represents the accumulation of a fluorescent binding fragment of tetanus neurotoxin (HCT) and a fluorescently conjugated antibody directed against the p75^NTR^ neurotrophin receptor (α-p75^NTR^) in ES-derived MNs following incubation with 2 μM of the different compounds. The yellow box shows threefold average standard deviation of the entire dataset. Compounds falling outside the yellow box in the upper right quadrant significantly enhanced accumulation. A1, A4 ^4^, E4 ^22^ and C3 were identified as potential transport enhancers.**B**. Western blot of SH-SY5Y cell lysates following treatment with recombinant GDNF and GFRα1, the RET inhibitors (C3 and LOXO-292), and their vehicle control, DMSO. SH-SY5Y cells were pre-treated with DMSO, 1 µM or 2 µM C3 or LOXO-292 for 1 h, and stimulated with 100 ng/ml GDNF and GFRα1 for the final 10 min of incubation.

In this study, we have investigated one hit from the screen, the compound C3 (Fig. 1A), which inhibits RET, a tyrosine kinase receptor for the glial cell-line derived neurotrophic factor (GDNF) family of ligands, including GDNF itself. Together with its coreceptor, GDNF-family receptor (GFR) α1, GDNF triggers RET dimerization, and activation of downstream signalling cascades, including PI3K-AKT and MAPK-ERK1/2, which promote neuronal survival and proliferation^23^. Although aberrant GDNF-RET signalling occurs in several types of cancer^24,25^, physiological RET signalling is also fundamental for MN development and survival^26^.

Alterations in GDNF-RET signalling are associated with ALS pathology, with increased microglial RET immunoreactivity in SOD1^G93A^ mice^27^, and increased GDNF levels in ALS patient muscle biopsies^28^. Furthermore, one of only two FDA-approved ALS therapeutics, edaravone, has been shown to increase RET and GFRα1 protein levels, and increase AKT and ERK1/2 activation in MNs derived from ALS-patient iPSCs^29^. However, although GDNF-based therapies have shown some promise, they have so far failed to progress from pre-clinical testing or produce significant clinical improvements^30,31^, with some studies even demonstrating adverse effects^32^. This highlights a complex relationship between the GDNF-RET signalling axis and ALS. Altogether, these findings support further testing of the hypothesis suggested by the pilot kinase inhibitor screen^4^ that altering RET signalling may rescue axonal transport deficits in ALS.

In this work, we identified a new functional interaction between the RET-GDNF signalling axis and axonal transport of signalling endosomes, and revealed profound differences in RET activity between WT and ALS neurons. We showed that pharmacological inhibition and knockdown of RET significantly enhance signalling endosome speed in WT MNs, and rescue *in vivo* axonal transport deficits in the SOD1^G93A^ mouse model of ALS.

## Results

### A kinase inhibitor screen identified RET inhibitors as potential axonal transport enhancers

To identify potential axonal transport modulators, we screened a small molecule library of kinase inhibitors in ES-MNs^4^. The readout for the screen was somatic accumulation of two fluorescent markers of signalling endosomes: the non-toxic heavy chain fragment of tetanus neurotoxin, conjugated to AlexaFluor555 (HCT-555); and a fluorescently-conjugated antibody against the extracellular domain of p75^NTR^ (α-p75^NTR^)^8,9^. An increase in the somatic accumulation of these probes in ES-MNs reflects a rise in the retrograde transport flow. Compounds that significantly increased the somatic accumulation of these retrograde probes fall outside of the yellow box in the upper right quadrant, which represents threefold average standard deviations of the entire dataset (Fig. 1A). Previous validation of p38MAPK (A1 and A4) and IGF1R (E4) inhibitors has revealed that these molecules are able to rescue retrograde transport deficits *in vivo* in symptomatic SOD1^G93A^ mice^4,22^.

Another hit from the screen was the RET inhibitor C3 (GW440139A; red box, Fig. 1A). Although initial investigation confirmed that 10 μM C3 significantly enhanced *in vitro* signalling endosome transport (data not shown), this concentration also induced neuronal blebbing (Supplementary Figure 1A). Using C3 at 2 μM stopped axonal blebbing, but at the same time abolished any significant effect on axonal transport speeds (data not shown).

To circumvent this problem, we synthesized a new batch of C3 compound and first assessed RET activation in the lysates of differentiated SH-SY5Y cells, which express high levels of RET^33^. We examined RET activation following treatment with C3, as well as LOXO-292, a commercially available RET inhibitor currently in clinical trials as an anti-cancer therapy^34^. As shown in Fig. 1B, both C3 and LOXO-292 inhibited RET activation to a similar extent (Fig. 1B and Supplementary Figure 1B), thus validating the new batch of C3. Importantly, at 1 μM, newly synthesized C3 did not cause significant neuronal blebbing in mixed ventral horn spinal cord cultures (hereafter referred to as primary MNs) in compartmentalized or mass cultures (Supplementary Figure 1C), suggesting that this phenomenon arose from impurities in the original C3 batch used in the kinase inhibitor screen (Fig. 1A).

### RET inhibition enhances axonal transport in primary WT motor neurons

To assess the effect of RET inhibition on axonal transport, we adapted a previously reported *in vitro* experimental set up^22^. As shown in Fig. 2A, primary MNs were cultured in compartmentalised microfluidic chambers (MFCs). On days *in vitro* (DIV) 6-7, both somatic and axonal compartments were treated with C3, LOXO-292 or vehicle control (DMSO), and 30 nM HCT-555 was added only to the axonal compartment for 45 min prior to live imaging. Axons were imaged at 2 frames/s by confocal microscopy, and transport dynamics were assessed through manual tracking with TrackMate^35^. Examples of kymographs of axons treated with vehicle control (DMSO) and RET inhibitors are shown in Fig 2B. C3 significantly increased average retrograde signalling endosome transport speeds in WT MNs (Fig. 2C, student’s t-test * p = 0.0395), and significantly reduced pausing (Fig. 2D, student’s t-test ** p = 0.0067). C3 did not influence maximum endosome speeds (Fig. 2E). Interestingly, the commercial RET inhibitor, LOXO-292, had no significant effect on axonal transport in WT MNs at 10 nM, 100 nM or 1 μM (Fig. 2F). Although non-significant, 100 nM LOXO-292 did induce a slight increase in WT endosome speeds (student’s t-test, p = 0.0845), suggesting that LOXO-292 may alter axonal transport speeds only when used at concentrations near to its IC50 (Ref 36).

**Figure 2.**
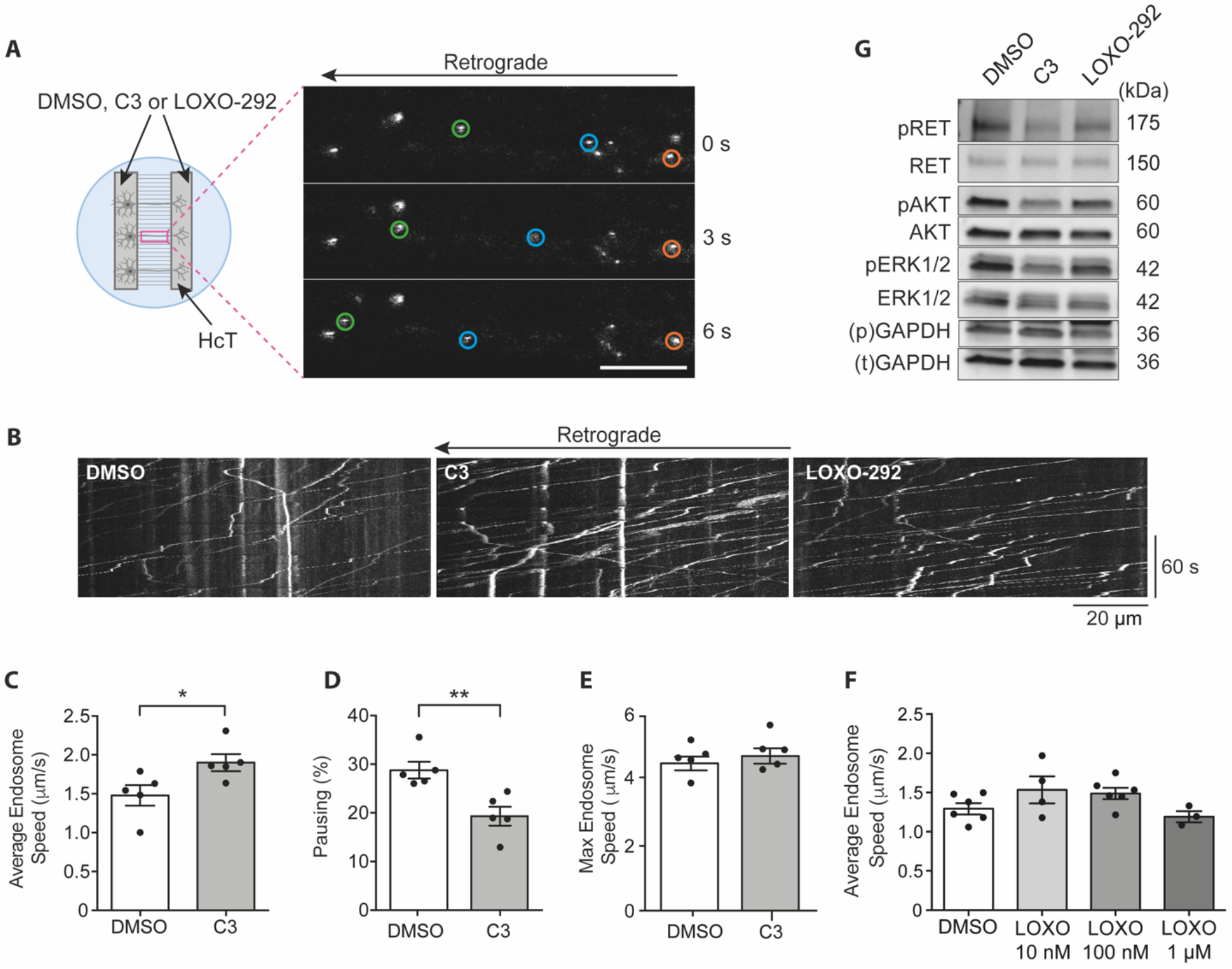
RET inhibition enhances axonal transport speeds in WT MNs. **A**. *In vitro* axonal transport experimental design. Mixed ventral horn cultures were seeded into the somatic compartment of 2-channel MFCs, and axons grew through the microgrooves and into the fluidically-isolated axonal compartment. DMSO, C3 or LOXO-292 were added to both compartments, whilst HCT was added only to the axonal compartment. Axons were imaged in the MFC microgrooves, and videos were captured at a frame rate of 2 frames/s. Videos were manually tracked using TrackMate to determine signalling endosome transport dynamics. Orange circles highlight a paused endosome, blue/green circles highlight retrogradely transporting endosomes. Scale bar, 20 µm. **B**. Representative kymographs of signalling endosome axonal transport videos in WT primary MN cultures following treatment with DMSO, 1 µM C3 or 100 nM LOXO-292. **C**. Average speeds of HCT-positive signalling endosomes following treatment with DMSO (empty bars) or 1 µM C3 (grey bars). C3 significantly increased mean signalling endosome speeds (* p = 0.0365, Student’s t-test). DMSO results were collected from the analysis of 414 endosomes in 31 videos for a total of 42 153 single endosomal movements. C3 results represent 456 endosomes in 27 axons with a total of 35 154 single endosomal movements. **D**. Bar graph showing percentage pausing with DMSO vs C3 treatment. C3 causes a significant decrease in pausing (** p = 0.0067, Student’s t-test). **E**. Mean maximum endosome speed seen in MN cultures treated either with C3 or DMSO (p = 0.4945, Student’s t-test). **F**. Mean signalling endosome transport speeds after primary MN cultures were treated with DMSO or increasing concentrations of LOXO-292. There was no statistically significant difference at any concentration (one-way ANOVA p = 0.1219). **G**. Western blot assessing activation of RET signalling in WT primary MN lysates following treatment with 1 µM C3, 100 nM LOXO-292 or DMSO for 1 h. Quantification is shown in Figure 5 B-D. Bands were normalised to corresponding GAPDH loading control (phospho-“p”, total “t”). Each point represents a biological replicate. DMSO vs. C3 n = 5 biological replicates, DMSO vs. LOXO-292 n = 4-6 biological replicates. Bars represent mean ± SEM.

We then assessed the effect of 1 μM C3 and 100 nM LOXO-292 on the activity of RET and its downstream signalling cascades. Interestingly, we found both C3 and LOXO-292 have a weaker inhibitory effect on RET activation in MNs compared to SH-SY5Y cells (Fig. 2G). In particular, in WT MNs, neither inhibitor inhibited RET autophosphorylation. However, C3 blocked the downstream activation of AKT and ERK1/2, two established components of the GDNF-RET signalling cascade in non neuronal cells^23^. Interestingly, no significant inhibition of AKT or ERK1/2 was seen using LOXO-292 (Fig. 2G, 5B) possibly explaining its weaker effect on signalling endosome transport compared to C3.

### GDNF stimulation enhances signalling endosome transport in WT motor neurons

We next assessed the effect of applying exogenous GDNF and GFRα1 on signalling endosome transport in WT MNs. To minimise variability between dishes, we cultured neurons in 3-channel MFCs, where axons project into two fluidically isolated compartments (Fig. 3A). This allowed the treatment of the same culture dish with both the vehicle control (PBS) and experimental condition (GDNF, GFRα1 or GDNF+GFRα1). GDNF stimulation (100 ng/ml) significantly increased signalling endosome transport speeds (Fig. 3B, E), whereas 100 ng/ml GFRα1 treatment alone had no effect (Fig. 3C, E). Since combining 100 ng/ml GDNF with 100 ng/ml GFRα1 (Fig. 3D, E) increased signalling endosome transport speeds similarly to GDNF alone (Fig. 3B, E), the enhancement in axonal transport is likely to be due to the application of GDNF, but not GFRα1, to primary MNs. Henceforth, GFRα1 is not rate limiting in this experimental system.

**Figure 3.**
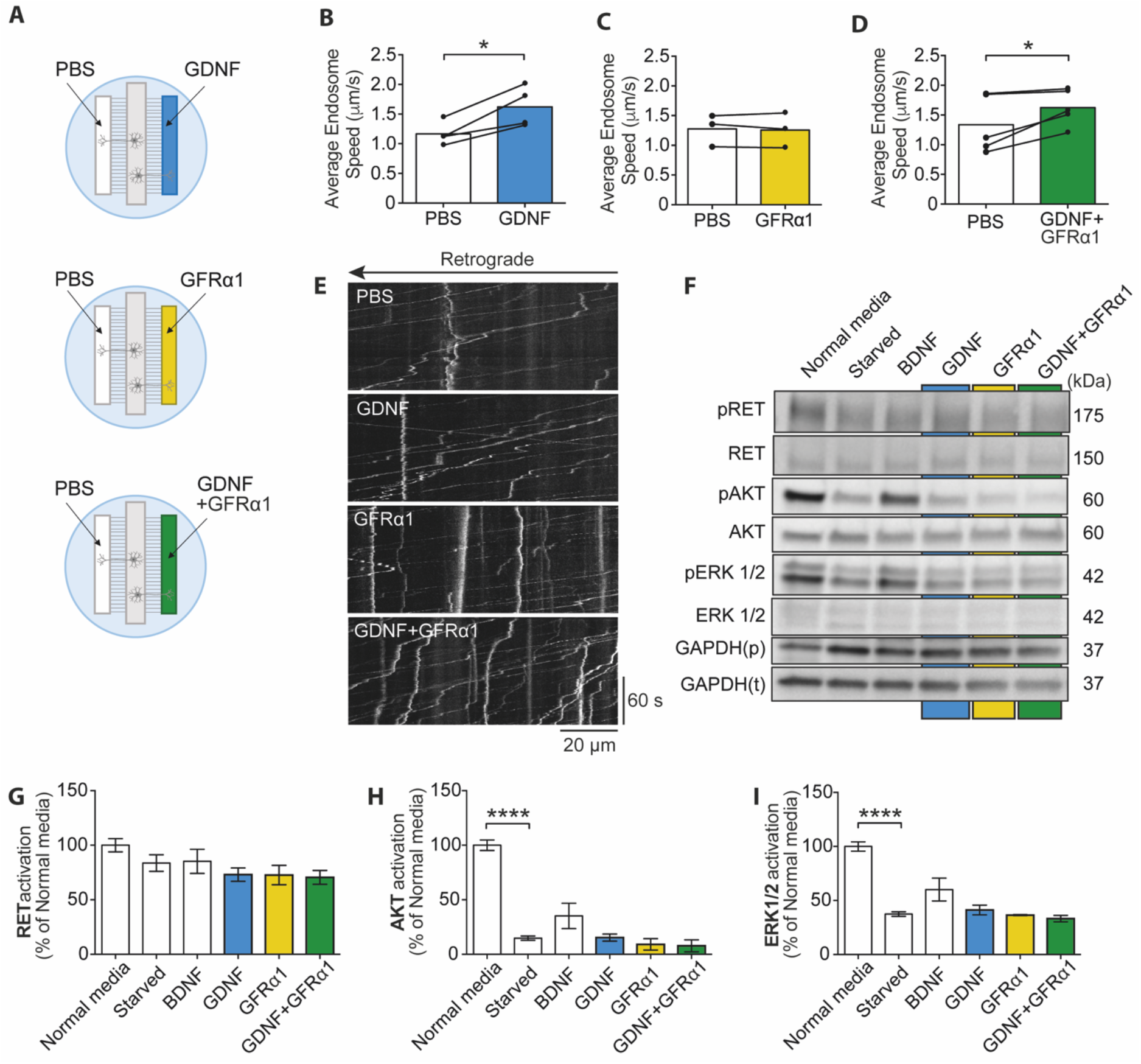
Stimulation with GDNF enhances WT axonal transport. **A**. Schematic representation of GDNF and GFRα1 transport experiments in three-compartment MFCs. Primary ventral horn neurons were cultured in the central (somatic) compartment of 3-channel MFCs, with axons projecting to two lateral fluidically isolated axonal chambers. 100 ng/ml GDNF, GFRα1, or both together were added to one axonal compartment, with PBS added to the other as an internal control. **B-D**. Average endosome speeds following the addition of GDNF (B), GFRα1 (C), and GDNF+GFRα1 (D) in the right compartment and PBS in the left compartment (B-D). Each point represents a biological replicate. Connecting lines represent speeds observed in control and experimental condition-treated axons from the same cultures. Speeds were significantly increased with GDNF alone (* p = 0.02) and GDNF with GFRα1 (* p = 0.04), but not GFRα1 alone (p = 0.6799), as determined by two-tailed paired t-tests. **E**. Kymographs showing signalling endosome transport in axons of WT primary MN cultures following treatment with PBS, GDNF, GFRα1 or GDNF together with GFRα1.N = 3-5 biological replicates. Bars represent mean ± SEM. PBS results were collected from 443 endosomes in 42 videos for a total of 45 569 single movements (n = 5). GDNF results were collected from 218 endosomes in 22 videos for a total of 17 938 single movements (n = 4). GFRα1 results were obtained from 216 endosomes in 24 videos for a total of 21 981 single movements (n = 3). GDNF and GFRα1 results were collected from 400 endosomes in 29 axons for a total of 31 809 single movements (n = 5). **F**. Western blot analysis of primary MN lysates following starvation in Neurobasal and Glutamax, and treatment with 50 ng/ml BDNF, or 100 ng/ml GDNF, and GDNF and GFRα1 (both 100 ng/ml). **G-I**. Western blot quantification of RET, AKT and ERK1/2 activation after starvation and stimulation with BDNF, GDNF and GFRα1. Starvation significantly decreased activation of AKT (**** p < 0.0001) and ERK (**** p < 0.0001), but not RET (one-way ANOVA with Dunnett’s multiple comparison test).

Then, we investigated the downstream signalling output in starved WT MNs following GDNF treatment in the presence or absence of exogenous GFRα1. Interestingly, we observed no increase in RET, AKT or ERK1/2 activation (Fig. 3F-I). The same cultures were, however, responsive to BDNF, which also induces activation of AKT and ERK1/2 via activation of TrkB^37^. Upon BDNF treatment (10 min, 50 ng/ml) activation of AKT and ERK1/2 increased above starvation levels, whereas there was no comparable increase with GDNF+/-GFRα1 treatment (Fig. 3F-I). Interestingly, we have previously shown that stimulation with this concentration of BDNF enhances axonal transport in WT primary MNs^7^. This implies that the enhancement of signalling endosome transport upon GDNF treatment is likely to be independent of RET activation, suggesting GDNF may activate an alternative receptor in these cells. These findings highlight a divergence in the signalling response to GDNF in MNs compared to cancer cell lines (e.g., SH-SY5Y cells), which are commonly used to investigate RET signalling^38,39^.

### Genetic downregulation of RET mimics the effect of RET inhibition on axonal transport

The negligible effect of LOXO-292 on signalling endosome transport, together with the significant enhancement observed with GDNF stimulation raises the possibility that the effect of C3 on axonal transport may result from off-target effects of this inhibitor. To confirm that reduction of RET signalling enhances retrograde transport, we assessed transport dynamics following lentiviral-mediated shRNA knockdown of RET in MNs.

Knockdown of RET was first established in mouse N2A neuroblastoma cells, where we selected the two most effective shRNAs to insert in lentiviral vectors (Supplementary Figure 2A-B). We then determined the optimal viral titre for knockdown in primary MN cultures (Supplementary Figure 2C). Lentiviral expression of both shRNAs induced robust RET knockdown in primary MNs in mass culture (Fig. 4A-C). These RET-targeting shRNAs increased the average (Fig. 4D, E) and maximum (Fig. 4D, F) signalling endosome transport speeds compared to scrambled control, without significantly altering the pausing of the organelles (Figure 4G), as seen using the C3 inhibitor (Fig. 2D). These findings validate C3 as specific RET inhibitor and support the role of this class of compounds as retrograde axonal transport enhancers.

**Figure 4.**
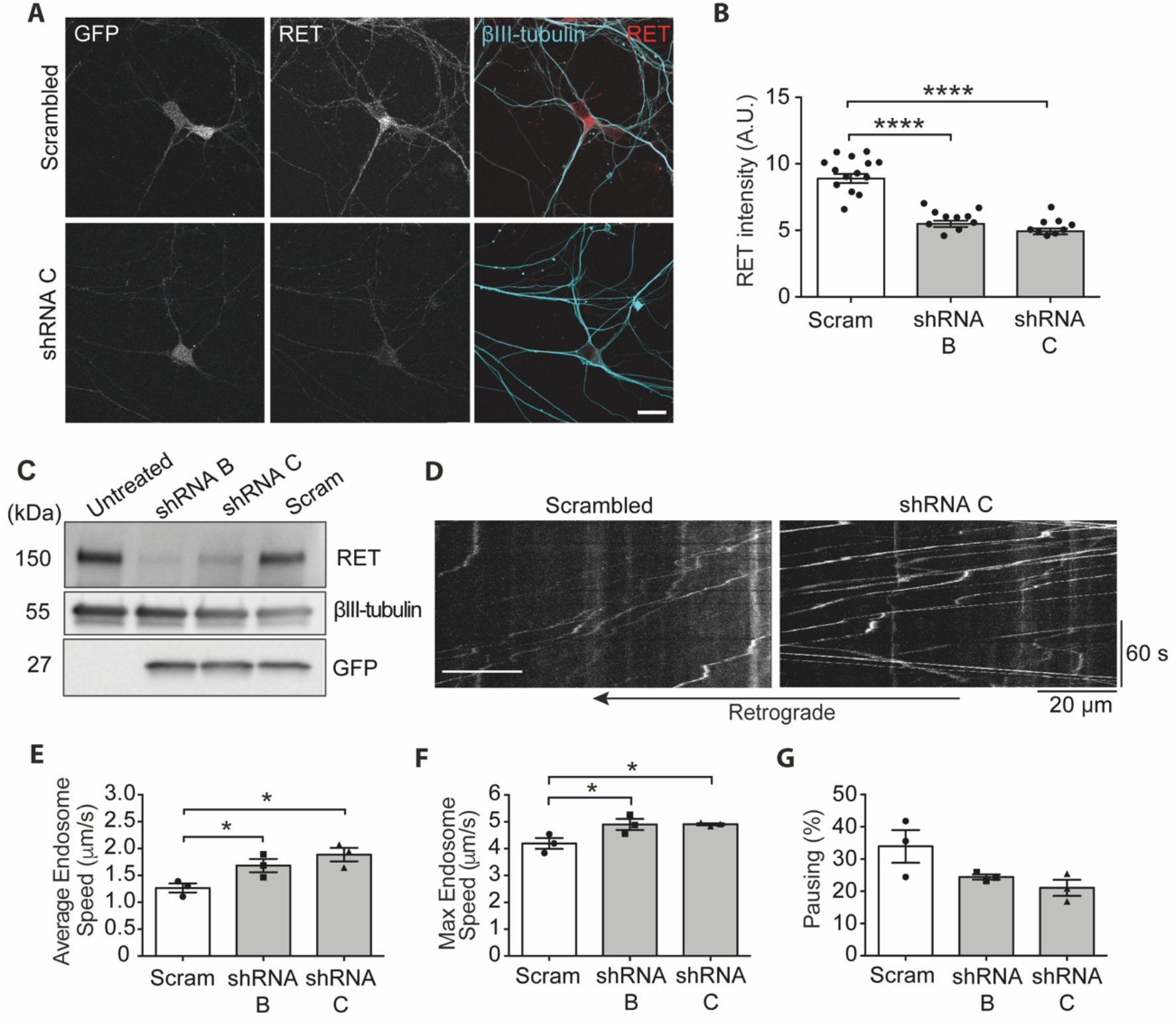
RET knockdown mimics the effect of pharmacological RET inhibition on axonal transport. **A**. Representative images of RET, βIII-tubulin and GFP staining in primary MN cultures transduced with scrambled control (Scram) or RET-targeting shRNA (shRNA C). Scale bar, 20 µm. **B**. Quantification of RET signal using a βIII-tubulin mask, as shown in A. **** p<0.0001 and determined by a one-way ANOVA. **C**. RET, GFP and βIII-tubulin levels in primary MN lysates after transduction with lentiviral particles expressing RET-targeting shRNA B, C and scrambled control (Scram). **D**. Representative kymographs from RET knockdown transport experiments. **E-G**. Primary ventral horn neurons after RET knockdown showing the E) mean signalling endosome transport speed (p<0.05 one-way ANOVA with multiple comparisons), F) maximum endosome track speed, and G) percentage pausing (p<0.05 one-way ANOVA with multiple comparisons). N = 3 biological replicates, bars represent mean ± SEM. Scrambled results were collected from 172 endosomes in 19 videos for a total of 16 870 single movements. The shRNA B dataset was collected from 162 endosomes in 17 videos for 14 145 single movements. The shRNA C dataset was collected from 107 endosomes in 11 videos for 8 080 single movements.

### RET signalling in the SOD1^G93A^ mouse model of ALS

Having shown that blockade of RET signalling enhances signalling endosome transport in WT neurons, we next investigated the effect of RET inhibition in SOD1^G93A^ MNs. We observed no significant difference in the effect of C3 and LOXO-292 on AKT or ERK1/2 activation in WT and SOD1^G93A^ neurons (Figs. 5A, C-D). However, RET activation levels in SOD1^G93A^ MNs were significantly reduced compared to WT (Figs. 5A-B, complete representative WT blot shown in Figure 2G). This evidence suggests the presence of altered RET signalling in SOD1^G93A^ neurons, warranting further investigations of RET signalling in ALS.

**Figure 5.**
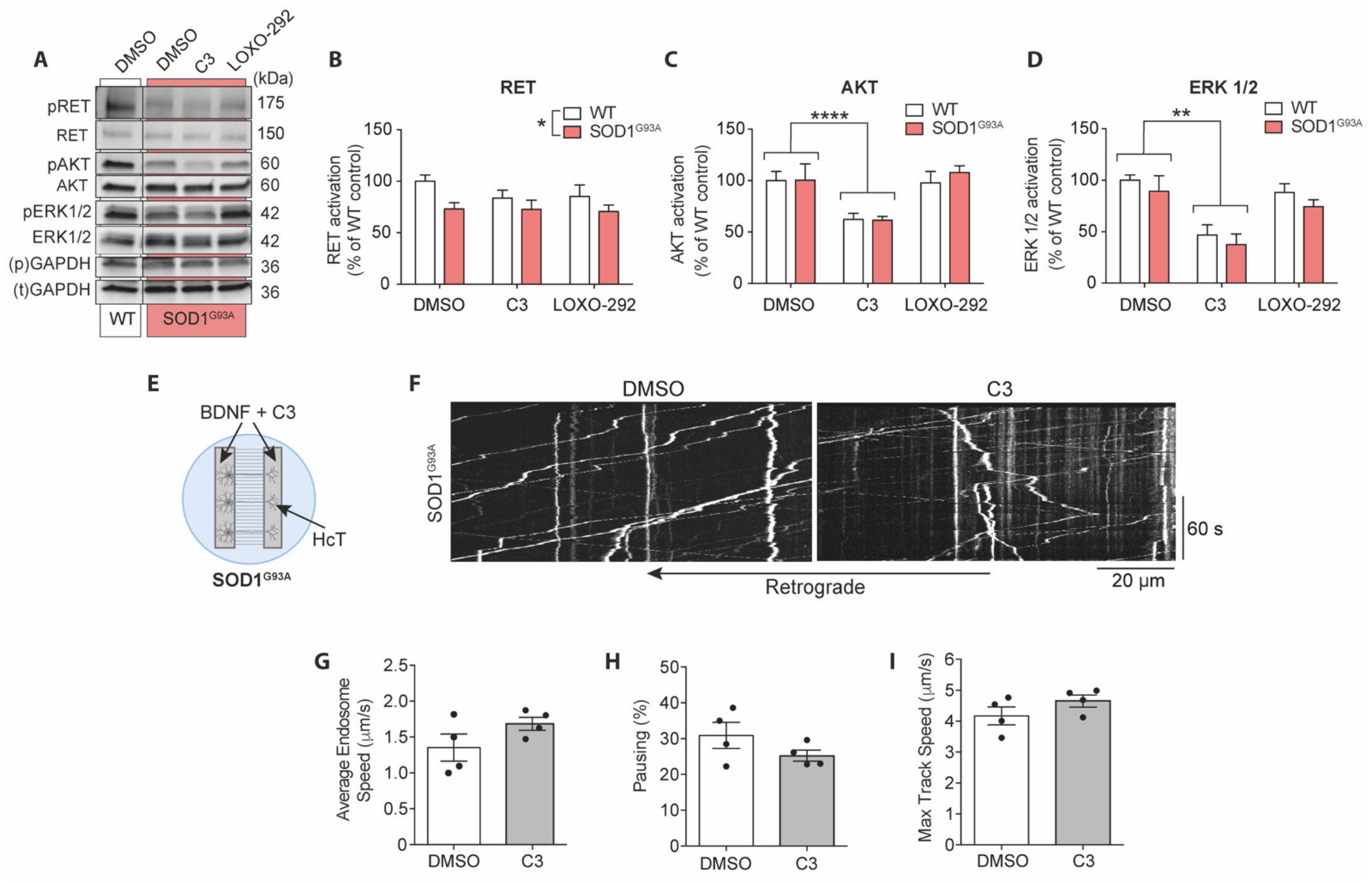
RET inhibition differentially impacts SOD1^G93A^ MNs compared to WT. **A**. Western blot showing WT and SOD1^G93A^ primary MN lysates (mass culture) following treatment with DMSO, 1 µM C3 and 100 nM LOXO-292 for 1 h. Full WT blot is shown in Figure 2G. Samples were run twice on two gels to analyse phosphorylated and total proteins of the same molecular weight. Bands were normalised to corresponding GAPDH loading control (phospho-“p”, total “t”). **B**. Quantification of RET activation from western blots of WT and SOD1^G93A^ primary MN lysates (1 µM C3, 100 nM LOXO-292; see also Fig. 2). There was no significant difference in RET activation comparing DMSO to either RET inhibitor (two-way ANOVA p = 0.1629); however, SOD1^G93A^ cultures have reduced RET activation compared to WT (two-way ANOVA; * p = 0.0181). **C**. Quantification of AKT activation. C3 significantly inhibits AKT activation compared to DMSO (two-way ANOVA with Tukey’s multiple comparison test; **** p<0.0001), but LOXO-292 has no significant effect. **D**. Quantification of ERK1/2 activation. C3 significantly inhibits ERK1/2 activation compared to DMSO (two-way ANOVA with Tukey’s multiple comparison test; ** p = 0.0018). In contrast, LOXO-292 has no significant effect. RET, AKT and ERK1/2 activation was determined by first normalising all bands to GAPDH to control for loading, then normalising phosphoprotein levels to total protein. Bars represent mean ± SEM. **E**. Experimental set up for SOD1^G93A^ MNs transport analysis following treatment with the C3 inhibitor. 50 ng/ml BDNF was added to both MFC chambers, whereas HCT was added to the axonal compartment only. **F**. Representative kymographs from axonal transport videos in SOD1^G93A^ primary MN cultures. Scale bars, 20 µm. **G-I**. Average endosome speeds, p = 0.1646 (G), percentage of pausing, p = 0.2002 (H) and maximum endosome speed, p = 0.2175 (I) in SOD1^G93A^ primary MN cultures following DMSO or 1 µM C3 treatment. Statistics determined by student’s t-tests. N = 4 biological replicates. Bars represent mean ± SEM. DMSO results were collected from 295 endosomes in 26 axons for a total of 29 667 single endosomal movements. C3 results were collected from 354 endosomes in 23 axons for a total of 30 664 single endosomal movements.

We next assessed signalling endosome transport dynamics in SOD1^G93A^ primary MNs. The experimental set up is shown in Fig. 5E. Neurons were treated with 1 μM C3 in the presence of 50 ng/ml BDNF to determine whether RET inhibition rescues the BDNF-dependent retrograde transport deficits previously described in SOD1^G93A^ primary MNs^5,7^. C3 treatment had no significant effect on average or maximum endosome speeds, although there is a trend for an increase in both, as well as a reduction of pausing (Fig. 5F-I).

### RET inhibition rescues retrograde transport deficits in the SOD1^G93A^ mouse model of ALS

To independently assess the effect of RET inhibition on axonal transport, we tested the C3 inhibitor *in vivo* by measuring the rate of transport of signalling endosomes in the intact sciatic nerve of early symptomatic (∼P73) SOD1^G93A^ mice, a time point that corresponds with ∼20% loss of lumbar motor neurons^9^. Using a previously optimised experimental design^40,41^, HCT-555 and 25 ng BDNF were injected into the tibialis anterior and gastrocnemius muscles in the lower limb of anaesthetised female WT and SOD1^G93A^ mice. For *in vivo* delivery, we suspended C3 in 1 % methyl cellulose (MC), and injected it intraperitoneally at a final concentration of 10 mg/kg. Since no pharmacokinetic or pharmacodynamic data for C3 is currently available, we selected 10 mg/kg based on previous findings showing that two p38MAPK inhibitors administered at both 10 mg/kg and 100 mg/kg significantly enhanced signalling endosome transport in SOD1^G93A^ mice *in vivo*^4^. Approximately 4 hours after injection of HCT-555, mice were re-anaesthetised and the sciatic nerve of the injected limb was exposed for intravital imaging.

Stills from representative transport videos are shown in Figure 6A. In SOD1^G93A^ mice treated with vehicle (1% MC in PBS), we observed the previously reported deficit in signalling endosome transport (Fig. 6B and Supplementary Figure 3A, B)^4,9,22^. Interestingly, we saw a significant increase in average endosome speed upon treatment of SOD1^G93A^, but not WT, mice with 10 mg/kg C3 (Fig. 6B). Accordingly, C3 induced a significant reduction in pausing in SOD1^G93A^ mice, without any significant effect in WT animals (Fig. 6C). We also show a slight shift to the right of the speed distribution curve of SOD1^G93A^ mice treated with the C3 inhibitor (Supplementary Figure 3D), indicating a general increase in axonal transport speeds upon RET inhibition. Surprisingly, however, C3 had no statistically significant effect on maximum endosome speed in WT or SOD1^G93A^ sciatic nerves (Fig. 6D). Although there was no difference in average endosome speeds between control and C3-treated WT mice, we did observe a slight shift to the left of the speed distribution curve following C3 treatment (Supplementary Figure 3C), which suggests that in WT mice, RET inhibition may trigger a subtle slowing down of axonal transport *in vivo*. Interestingly, we found that GDNF has no effect on WT or SOD1^G93A^ signalling endosome transport speeds *in vivo* (Supp. Fig. 4A-C), indicating that *in vivo*, increasing GDNF signalling has no impact on axonal transport, and RET inhibition only enhances transport speeds where GDNF-RET signalling is pathologically altered. These results further support dampening rather than boosting GDNF-signalling for the treatment of ALS.

**Figure 6.**
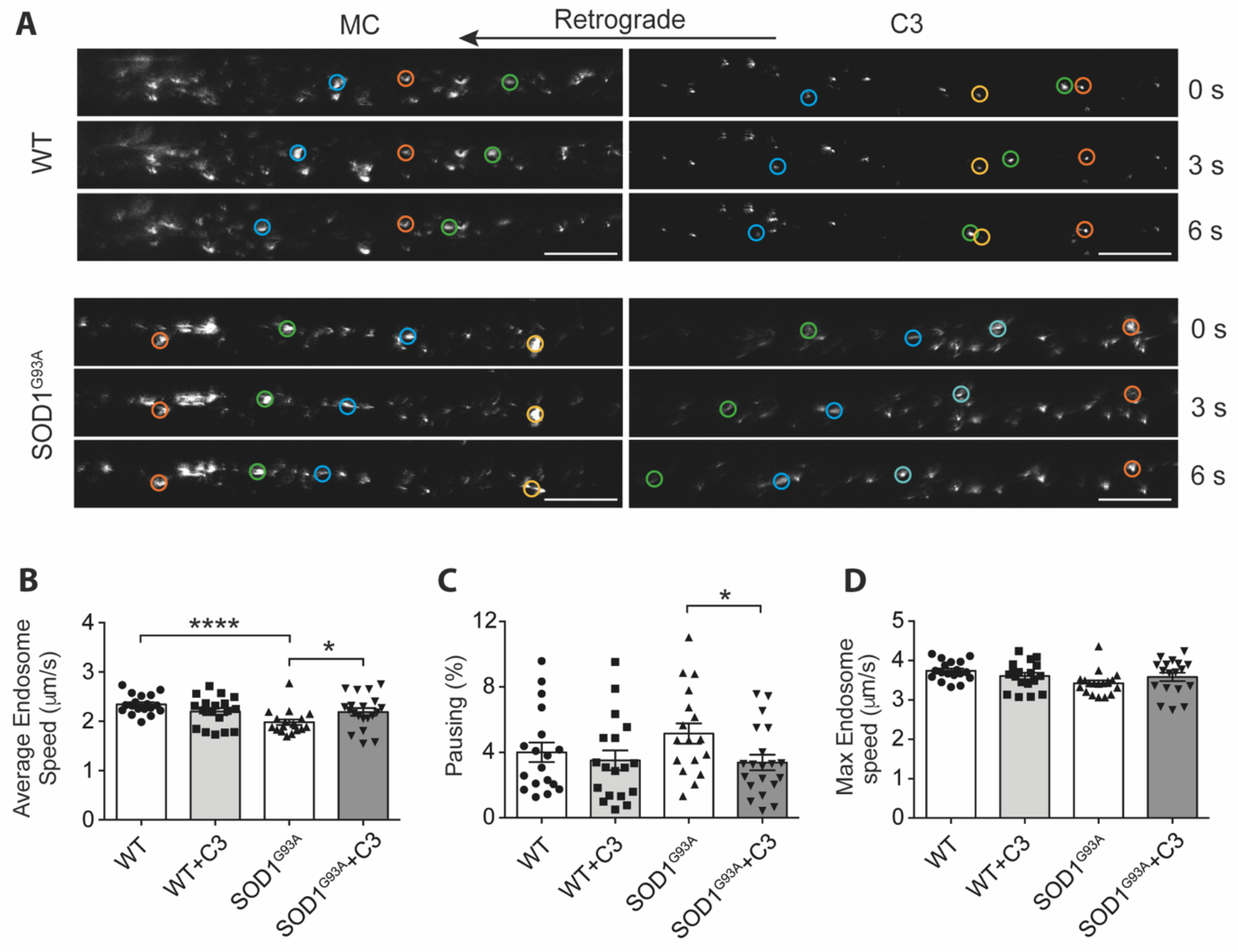
RET inhibition increases *in vivo* transport speeds in the SOD1^G93A^ mouse model of ALS. **A**. Still frames from signalling endosome transport videos in the sciatic nerve of WT and SOD1^G93A^ mice after intraperitoneal injection of 10 mg/kg C3 or vehicle control (1% methyl cellulose). Coloured circles highlight frame-to-frame endosomal movements (orange/yellow represent paused endosomes, blue/green represent retrogradely transporting endosomes). Scale bars, 20 µm. **B-D**. Average speeds (B), percentage pausing (C) and maximum transport speeds (D) of *in vivo* signalling endosome transport in the sciatic nerve of P73 SOD1^G93A^ mice following C3 treatment compared to vehicle control (MC; 1% methyl cellulose in PBS). C: **** p <0.0001; * p = 0.0415; D: * p = 0.0284; Student’s t-test. N = 6 animals. Statistics carried out per axon (3 per mouse, 18 per condition). Bars represent mean ± SEM.

Altogether, these results suggest that RET inhibition has the potential to rescue retrograde transport deficits in the SOD1^G93A^ mouse model of ALS. Optimisation of *in vivo* dose and route of delivery of C3 may potentiate the effect observed, promoting RET inhibition as a possible therapeutic approach in the treatment of ALS.

## Discussion

Deficits in the retrograde axonal transport of signalling endosomes and other organelles are one of the earliest changes observed in several models of ALS^6,9^. Protein kinases are known to influence axonal transport^15^, and we have previously shown that inhibition of both IGF1R and p38MAPK is effective in rescuing these deficits ^4,22^. The literature investigating GDNF-RET signalling predominantly originates from experiments in cancer cell lines, such as SH-SY5Y neuroblastoma lines, which express RET at high levels following retinoic acid differentiation^38^. Whereas this is informative to define the mechanism by which RET mutations impact cancer initiation and progression^23^, a comprehensive analysis of RET signalling in neurons is needed to further our understanding on the implications of deregulation of RET signalling in neuromuscular diseases. In this study, we explored the role of the GDNF-RET signalling axis on axonal transport in MNs and found that both GDNF stimulation and RET inhibition boost axonal transport of signalling endosomes. We have validated RET inhibitors as axonal transport enhancers with the potential to rescue disease-dependent axonal transport deficits *in vivo*, and revealed a fundamental function of GDNF-RET signalling in the axonal transport dynamics of signalling endosomes.

Reducing RET activation, either through pharmacological or genetic approaches, increases retrograde axonal transport dynamics (Fig. 2, 4 and 6). Despite inducing significant increases in axonal transport speeds in WT neurons *in vitro*, C3 had no significant effect on *in vivo* transport speeds in WT mice at the concentration investigated in this study. Interestingly, we also saw no increase in signalling endosome transport speeds *in vivo* following GDNF treatment in WT or SOD1^G93A^ mice. We showed that RET inhibition in SOD1^G93A^ primary MNs is not sufficient to completely overcome the transport impairment observed *in vitro*. The increased variability of endosome speeds in control conditions (DMSO) in these experiments is the likely determining factor preventing treatment with C3 from reaching significance in SOD1^G93A^ MNs. Furthermore, the co-treatment with BDNF might further mask the effect of RET inhibition on axonal transport. In fact, several signalling pathways downstream of RET are activated upon BDNF stimulation, such as AKT and ERK1/2^23,37^. Therefore, activation of TrkB by BDNF may hide any positive effect on transport induced by RET inhibition. Furthermore, the significant reduction in baseline RET signalling in SOD1^G93A^ MNs compared to WT observed *in vitro* could also contribute to the differential response to RET inhibitors of WT and SOD1^G93A^ neurons (Fig. 5B), providing an explanation as to why RET inhibition has a reduced impact in these neurons. Interestingly, however, acute treatment of symptomatic SOD1^G93A^ mice does rescue *in vivo* retrograde transport deficits monitored in the sciatic nerve. These findings highlight a divergence in the signalling response between primary MNs and the adult sciatic nerve, which could be attributed to the increased duration of C3 treatment in *in vivo* experiments (∼4 h *in vivo* vs. 1 h *in vitro*), as well as the developmental stage of the models.

Additionally, these findings suggest that *in vivo*, C3-mediated RET inhibition selectively targets pathologically altered RET signalling. Changes in GDNF and RET signalling have been previously reported in ALS, with increased immunoreactivity of RET in activated microglia, and reduced expression of GFR*α*1 mRNA in the spinal cord of SOD1^G93A^ mice^42,43^. Although edaravone, one of only two FDA-approved therapeutics for ALS, has been shown to increase RET, GDNF and GFR*α*1 protein levels^29^, enhancing GDNF signalling in ALS has so far either induced adverse effects ^32^ or failed to translate to significant clinical improvements^31^. These negative outcomes align to independent clinical trials in Parkinson’s disease^44^. Together with our results, these findings support the conclusion that dampening rather than enhancing RET signalling may have therapeutic promise in ALS.

Neither C3, nor the commercial RET inhibitor LOXO-292, significantly inhibited RET phosphorylation in primary MNs; however, C3 does halt AKT and ERK1/2 activation in WT and SOD1^G93A^ neurons (Figs. 5C, D). We have previously shown that direct inhibition of AKT is sufficient to increase signalling endosome transport speeds^22^, therefore it is possible that C3 enhances transport through indirect inhibition of AKT. Along the same lines, the failure of LOXO-292 to induce a significant enhancement of signalling endosome transport (Fig. 2F) could result from its inability to inhibit AKT activation in primary MNs (Fig. 5C).

RET was first identified as a proto-oncogene^45^, and activating-mutations drive neoplastic processes^39^. Notably, RET rearrangements occur in multiple endocrine neoplasia type 2B (MEN2B)^46^, emphasising the therapeutic importance of targeting this kinase receptor. Accordingly, many RET inhibitors have been developed and are currently being tested for safety and efficacy^47^ or are in clinical trials, including LOXO-292^34^. Repurposing these agents for use in neurodegeneration would provide an opportunity for significantly faster access of ALS patients to novel therapeutics.

However, a conclusion emerging from our studies is that GDNF-RET signalling in MNs is more complex than in cancer cell lines. Both GDNF and RET clearly play a fundamental role in MN development and survival^26,48,49^, as evidenced by the inclusion of GDNF in standard MN media. GDNF has a robust effect on axonal transport speeds in WT neurons *in vitro*, however, this appears to be independent of its canonical receptor RET. GDNF stimulation of primary MNs fails to activate AKT and ERK1/2 (Fig. 3F), two key signalling proteins switched on downstream of GDNF-RET activation in several cell lines^50^. GDNF stimulation of MNs also fails to increase RET phosphorylation, suggesting that in these neurons, GDNF enhances WT signalling endosome transport through a RET-independent mechanism.

Interestingly, GDNF-GFRα1 can signal independently of RET. For example, GDNF induces axonal outgrowth in hippocampal neurons, which do not express RET^51^. This response occurs alongside an increase in fibroblast growth factor receptor activation, a kinase receptor activated downstream of neural cell adhesion molecule (NCAM)^52^. Furthermore, GDNF-mediated chemoattraction of Schwann cells, which express GFRα1 but not RET, was abolished in the presence of NCAM-blocking antibodies^53^. This effect required downstream activation of the tyrosine kinase, Fyn^54^. Interestingly, Fyn has been shown to promote anterograde transport through phosphorylation of the microtubule associated protein tau^55^, releasing its inhibitory effect on kinesin^56,57^. Furthermore, dorsal root ganglion neurons from Fyn^-/-^ knockout mice show impairments in both anterograde and retrograde transport^55^. Therefore, it is possible that stimulation of MNs with GDNF potentiates retrograde signalling endosome transport through activation of NCAM signalling.

These findings highlight a divergence of RET and GDNF signalling in MNs, promoting both enhancement of GDNF signalling and RET inhibition as possible therapeutic approaches, and perhaps explaining the uncertain outcomes of GDNF-based therapies. Further investigation of both GDNF and RET signalling cascades is required to determine the physiological outcomes of manipulating these pathways in neurons, which would enable new translational efforts in ALS and other neurodegenerative conditions.

## Acknowledgements

We thank James Dick and the personnel of the Denny Brown Laboratories for assistance in maintaining the mouse colonies (Queen Square Institute of Neurology, University College London), and James N. Sleigh (Queen Square Institute of Neurology, University College London) for critical reading of the manuscript. This work was supported by a Medical Research Council PhD Studentship (ERR); a Junior Non-Clinical Fellowship from the Motor Neuron Disease Association (Tosolini/Oct20/973-799) (APT); a Leonard Wolfson PhD studentship (ADF); the ARUK UCL Drug Discovery Institute is core funded by Alzheimer’s Research UK (520909) (WM), Wellcome Senior Investigator Awards (107116/Z/15/Z and 223022/Z/21/Z) (GS), and a UK Dementia Research Institute Foundation award (UKDRI-1005) (GS).

## Author Contributions

Conceptualisation: ERR and GS. Investigation: ERR, APT, and ADF. Provision of essential reagents: WM. Writing: ERR and GS, with input from all authors. Funding: GS. All authors approved this submission. The funders had no role in study design, data collection and analysis, decision to publish, or manuscript preparation.

## Materials and Methods Animals and tissue collection

All experiments were conducted under the guidelines of the UCL Institute of Genetic Manipulation and Ethics Committees, and in accordance with the European Community Council Directive of 24 November 1986 (86/609/EEC). Animal experiments were performed under license from the UK Home Office in accordance with the Animals (Scientific Procedures) Act 1986 and were approved by the UCL Institute of Neurology Ethical Review Committee. Transgenic mice heterozygous for the mutant human SOD1 (G93A) on a C57BL/6-SJL mixed background (B6SJLTg [SOD1*G93A]1Gur/J) were acquired from The Jackson Laboratory. Colonies were maintained at the Queen Square Institute of Neurology Biological Services Unit. Heterozygous males were mated with wild-type C57BL/6-SJL F1 generation females to obtain experimental colonies. Mouse genotypes were confirmed using ear biopsies and PCR amplification of the human SOD1^G93A^ transgene as described by The Jackson Laboratory. Animals were housed in a controlled temperature and humidity environment and maintained on a 12 h light/dark cycle with access to food and water provided ad libitum.

### Plasmids and reagents

Chemicals were from Sigma-Aldrich unless otherwise stated. Compound C3 (GW440139A), originally provided by GSK (UK), was re-synthesised at the ARUK Drug Discovery Institute at UCL^58^. LOXO-292 was purchased from MedChemExpress (USA). Recombinant human GDNF and BDNF were purchased from PeproTech (UK), and GFRα1 was purchased from R&D Systems (USA). RET shRNA constructs were purchased from Genecopoeia in psi-LVRU6GP plasmids with eGreen Fluorescent Protein (GFP) reporter genes and puromycin stable selection markers (MSH025822-LVRU6GP, USA). Antibodies used in immunofluorescence (IF) and western blot (WB) experiments were as follows: RET (C31B4, WB 1:1 000, IF 1:250), RET pY905 (3221 1:1 000), ERK1/2 (9102 1:1 000), pERK1/2 T202/Y204 (9101 1:1 000), AKT (9272 1:1 000), pAKT S473 (4060 1:1 000, Cell Signalling Technology (UK); GAPDH (mab374 1:5 000, Merck Millipore, Germany); βIII-tubulin (IF 1:500, WB 1:3 000, 302305, Synaptic Systems, Germany), GFP (GFP-1010, IF 1:1 000, WB 1:5 000, Aves Labs, USA). The α-p75^NTR^ used in the kinase inhibitor screen was previously generated and characterised by the authors^8,59,60^.

### Cell line culture

SH-SY5Y, N2A and Lenti-HEK-293 cells were all cultured in Dulbecco’s Modified Eagle Medium (DMEM) with 10% fetal bovine serum (FBS). Cells were split every 2-3 days, once they reached 80-90% confluency. For experiments, cells were plated onto PDL-coated coverslips or straight onto plastic, cultured for 1 DIV, and either left undifferentiated (HEK-293), or differentiated using 10 μM retinoic acid (Sigma) for 12-48 h (SH-SY5Y and N2A).

### Motor neuron cultures

HB9-GFP ES-derived MNs were cultured as previously described^61^. GFP expression, driven by the HB9-homeobox gene enhancer allowed confirmation of MN identity. Mixed ventral horn cultures, referred to in this study as primary MNs, were isolated from E11.5-13.5 mouse embryos as previously described). Briefly, ventral horns from E11.5-13.5 SOD1^G93A^ and WT mice were dissociated, centrifuged at 1 500 rpm for 5 min, seeded into microfluidic chambers^62^, coverslips or wells, and maintained in MN media (Neurobasal (Gibco) with 2% v/v B27 (Gibco), 2% heat-inactivated horse serum, 1x GlutaMAX (Invitrogen), 24.8 µM β-mercaptoethanol, 10 ng/ml ciliary neurotrophic factor (CNTF; R&D Systems, USA) 0.1 ng/ml GDNF, 1 ng/ml BDNF and 1x penicillin streptomycin (Thermo Fisher)) at 37°C and 5% CO2.

### Lentiviral particle production and PMN transduction

RET shRNA and scrambled control viral particles were made by co-transfecting shRNA, packaging and envelope plasmids into Lenti-X 293 T cells (ClonTech, Mountain View, CA) with Lipofectamine 3000 (Thermo Fisher). Medium was collected at 48 h and 72 h after transfection. Medium containing lentiviral particles was concentrated using a Lenti-X concentrator (Takara, Japan) and resuspended in Opti-MEM (Thermo Fisher). shRNA viral particles were stored at -80°C until needed. Neurons were transduced on DIV3 by adding viral particles directly to the medium. Viral particles remained for 72 h, and neurons were lysed or assessed for transport on DIV6.

### *In vitro* retrograde transport assay

At 6-7 DIV, 30 nM AlexaFluor555-labelled HCT was added to the axonal compartmen ^22^, with key reagents (BDNF for SOD1^G93A^ primary MN experiments, GDNF, GFRα1, DMSO, LOXO-292, C3) or vehicle control (PBS) added to both axonal and somatic compartments for 45 min. Media was then replaced with fresh MN media containing 20 mM HEPES-NaOH (pH 7.4) and experimental compound for live imaging on an inverted Zeiss LSM 780 confocal microscope using a 40x oil immersed objective. Microscope and stage were pre-warmed to 37°C. Videos were taken at 0.500 s frame rate for >2.5 min using a resolution of 1024 × 1024 pixels.

### *In vivo* retrograde transport assay

The *in vivo* axonal transport was performed as previously described^40,41^. Briefly, P73 female WT (non-transgenic) and SOD1^G93A^ mice were anesthetized using isoflurane (National Veterinary Services) and AlexaFluor555-conjugated HCT (13 μg) and BDNF or GDNF (25 ng) were injected intramuscularly into the exposed tibialis anterior and lateral gastrocnemius muscles of the right hind leg and the wounds then sutured (for GDNF experiments, only tibialis anterior was injected). At this time, for C3 vs MC experiments only, mice were also intraperitoneally injected with either 10 mg/kg C3 or an equivalent volume of vehicle control (1% methyl cellulose (MC) [Sigma 274429]). Mice were then left to recover and kept under standard conditions with unlimited food and water supply. After 4+ h, mice were re-anesthetized using isoflurane and the sciatic nerve of the right leg was exposed. The anaesthetised mouse was placed on a heated stage in an environmental chamber and axonal transport was imaged in the intact sciatic nerve by time-lapse confocal microscopy as previously described ^40,41,63^. An inverted Zeiss LSM 780 with 40x Plan-Apochromat oil immersion objective was used to locate the sciatic nerve and visualise axons containing HCT-555 labelled signalling endosomes. Time lapse images were then acquired every 3 s using a resolution of 1024 × 1024 pixels. Following imaging, mice were immediately culled by cervical dislocation.

### Immunofluorescence

Cells on coverslips or in MFCs were fixed using 4% paraformaldehyde and 5% sucrose in PBS for 10 min then permeabilised in blocking solution (10% heat-inactivated horse serum in PBS, 0.5% BSA) with 0.2% Triton-X100. Permeabilised cells were incubated in primary antibody (detailed in Plasmids and Reagents) in block for 1 h at room temperature, washed five times in PBS, and then incubated in fluorescently-conjugated secondary antibodies (1:1 000) and DAPI (1:2 000) for 1 h at room temperature in the dark. Coverslips/MFCs were washed 5 times in PBS, then coverslips were mounted on glass slides using DAKO Mounting Medium, and imaged using an inverted Zeiss LSM 510 or 780 confocal microscope, on a 63x, 1.4 NA DIC Plan-Apochromat oil-immersion objective.

### Western blotting

Primary MN lysates were prepared in RIPA buffer (50 mM TrisHCl pH 7.5, 150 mM NaCl, 1% NP-40, 0.5% sodium deoxycholate, 0.1% sodium dodecylsulphate (SDS), 1 mM EDTA, 1 mM EGTA) with Halt™ protease and phosphatase inhibitor cocktail (1:100, Thermo Fisher), and incubated on ice for 30 min. Lysates were spun at 14 800 rpm, 4°C for 15 min, the supernatant collected, setting some aside to quantify protein concentrations. To the remaining lysate, 4x Laemmli sample buffer (15% SDS, 312.5 mM Tris-HCl pH 6.8, 50% glycerol, 10% β-mercaptoethanol, 0.1% bromophenol blue) was added, and samples were boiled for 5 min at 100°C. Pierce™ BCA protein Assay Kit (Thermo Fisher) was used to quantify the protein concentration. Protein separation was carried out using 4–15% Mini-PROTEAN® TGX Stain-Free™ protein gels (Bio-Rad, USA), and transferred onto 20% ethanol-soaked polyvinylidene difluoride (PVDF) membranes (Bio-Rad, USA). Membranes were blocked in 5 % BSA dissolved in Tris-buffered saline containing 0.1% Tween-20 (TBST) for 1 h at room temperature, then incubated with primary antibody diluted in 5 % BSA overnight at 4°C. Membranes were washed 4 × 5 min in TBST, then incubated with secondary antibody diluted in 5% BSA and washed again. Immunoreactivity was detected using chemiluminescent substrates (Luminata Crescendo/Classico, Merck Millipore) and the ChemiDoc™ Touch Imaging System with Bio-Rad ImageLab software. For blots where total and phosho-proteins of the same molecular weight were analysed, samples were run twice on two gels, and normalised to the corresponding GAPDH loading control (phospho “p” and total “t”). Original western blots are shown in Supplemental Material – Western Blot Originals.

### Image analysis

Image analysis was performed using Fiji (NIH, MD). For the analysis of signalling endosome transport dynamics, the Fiji plugin TrackMate^35^ was used to manually track cargo. TrackMate outputs “.csv” files, which provide information on the average and maximum speed of each endosome, as well as data describing single endosomal movements to determine instantaneous velocities and percentage pausing (frame-frame movements <0.2 µm/s). *In vitro*, all retrogradely moving endosomes were tracked. Due to the large volume of endosomes in *in vivo* videos, 25 endosomes were tracked per axon, in 3 axons per mouse. For both *in vitro* and *in vivo* experiments, endosomes pausing for >15 s were excluded from analyses. *In vitro* and *in vivo* speed distribution curves were generated by categorising instantaneous velocities into 0.2 µm/s bins.

## Statistical analysis

Statistical analysis was performed in Graphpad Prism (San Diego, California). Non-parametric Student’s t-test was used when comparing two groups. One-or Two-way analysis of variance (ANOVA), followed by Tukey’s or Holm-Šídák’s multiple comparisons tests were used when multiple groups were present. Significance is noted as follows: * p ≤ 0.05, ** p≤ 0.01, *** p ≤ 0.001, **** p ≤ 0.0001.

---

**Supplementary Figure 1.**
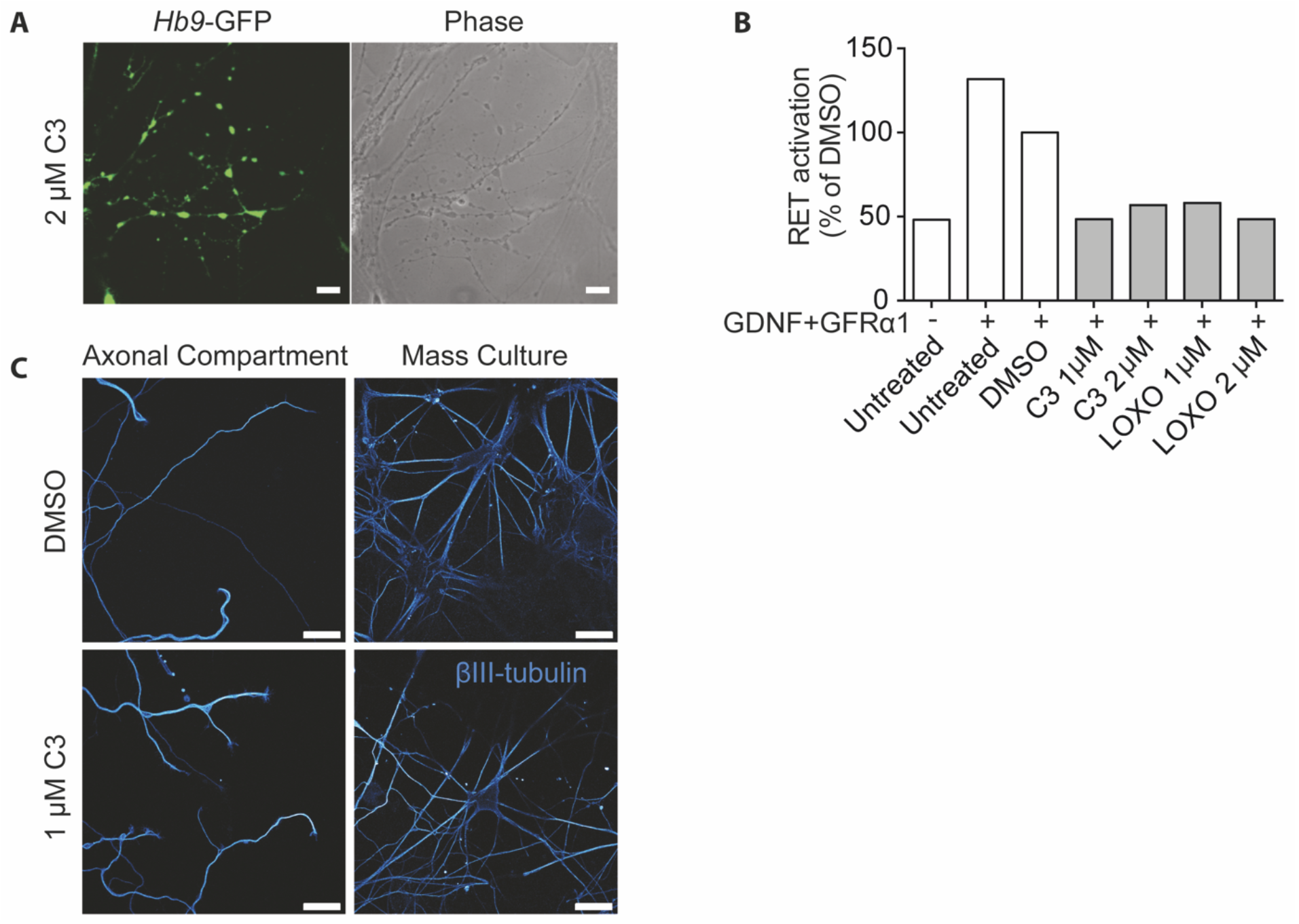
Newly synthesised C3 inhibitor does not cause neuronal blebbing. **A**. *Hb9*-GFP and phase contrast images of ES-derived MNs after treatment with 2 µM of the original C3 compound used in the kinase inhibitor screen. Neurons were imaged 3-5 days after embryoid body dissociation. Scale bar represents 10 µm. **B**. Quantification of RET activation after treatment with C3 and LOXO-292 observed in Fig.1B. RET activation was determined by normalising phospho RET band intensities to total RET, and is represented as a percentage of the average intensity of DMSO bands. **C**. βIII-tubulin staining of mixed ventral horn cultures in MFCs or mass culture following treatment with 1 µM of the newly synthesised batch of C3 or DMSO for 1 h. Scale bars, 20 µm.

**Supplementary Figure 2.**
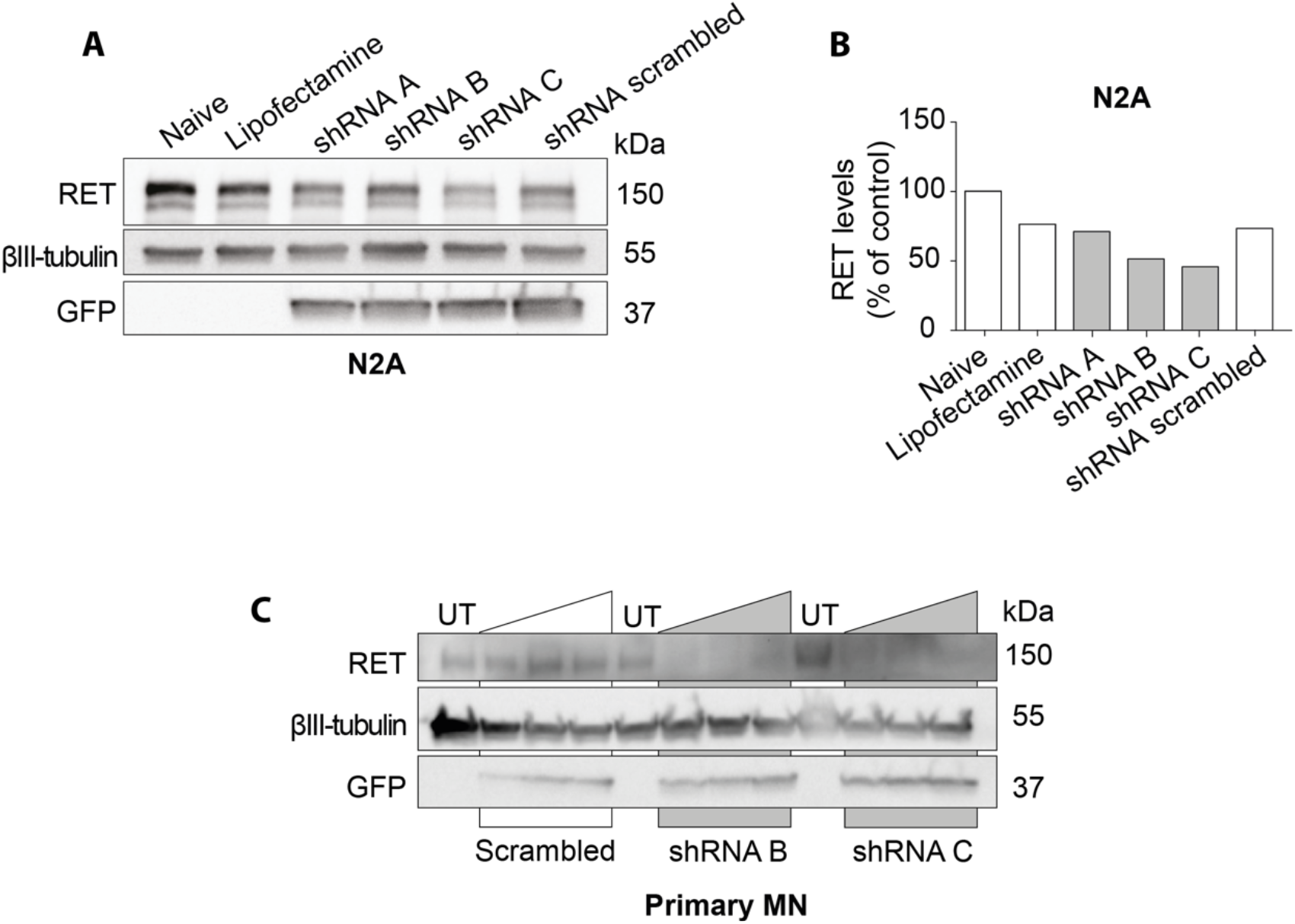
shRNAs targeting RET robustly reduce RET levels in N2A cells and primary MN cultures. **A**. RET, GFP and tubulin levels in N2A cell lysates after transfection with each RET-shRNA plasmid, and the scrambled control. **B**. Quantification of RET knockdown in N2A cells, relative to naïve sample. **C**. RET, GFP and tubulin levels in primary MN cultures after transduction with increasing volumes of lentiviral particles (2, 4 and 6 µl) expressing scrambled and RET-targeting shRNAs. N = 1 biological replicate for preliminary optimization.

**Supplementary Figure 3.**
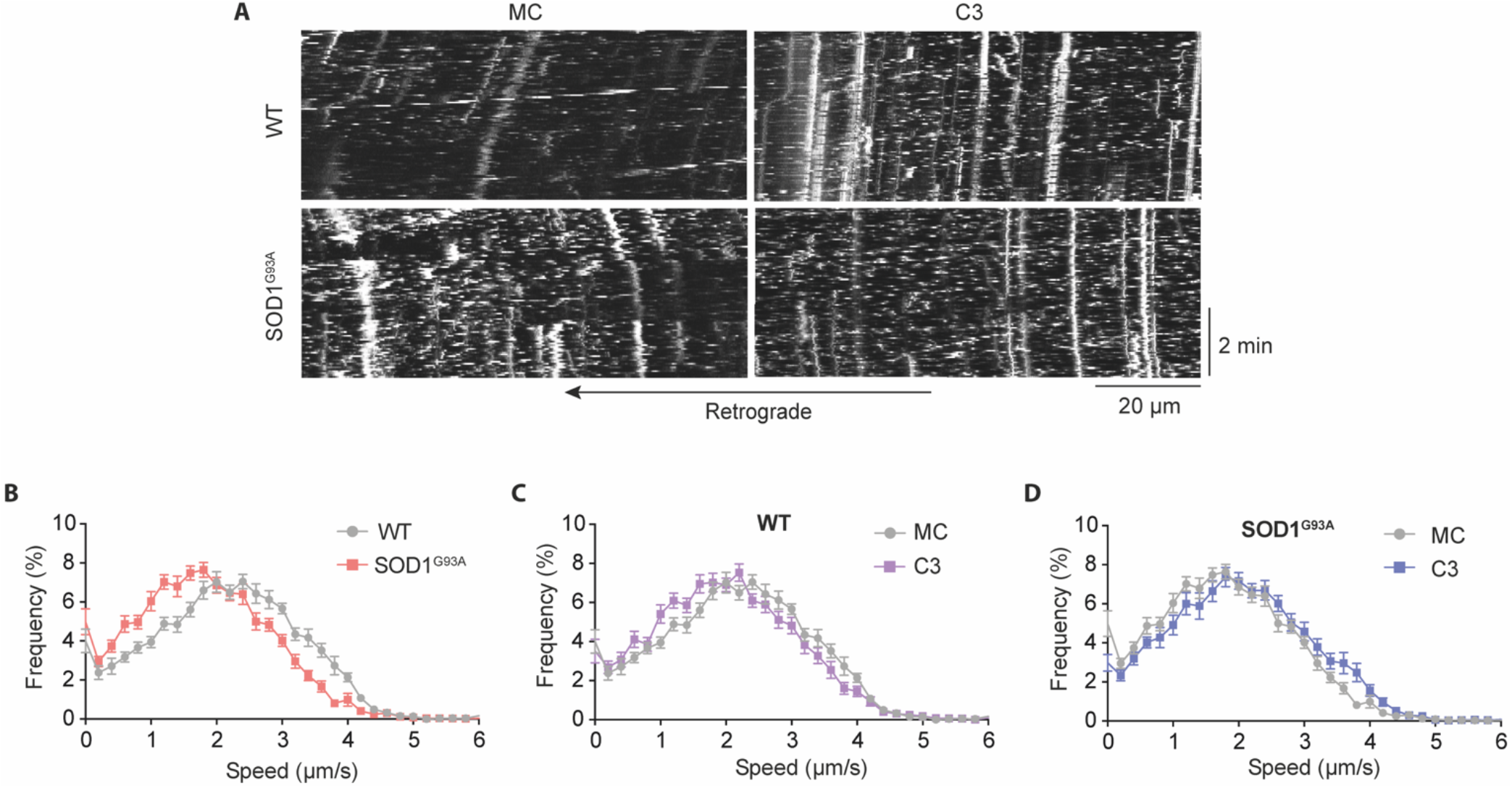
RET inhibition alters *in vivo* signalling endosome transport in the WT and SOD1^G93A^ sciatic nerves. **A**. Representative kymographs from axonal transport videos in WT and SOD1^G93A^ sciatic nerves. **B**. WT vs SOD1^G93A^ speed distribution curves after treatment with control vehicle (1% MC) vs C3 inhibitor. **C**. Speed distribution of signalling endosomes in WT sciatic nerve, comparing control vehicle vs C3 inhibitor. **D**. Speed distribution of signalling endosomes in SOD1^G93A^ sciatic nerve, after treatment with control vehicle vs C3 inhibitor.

**Supplementary Figure 4.**
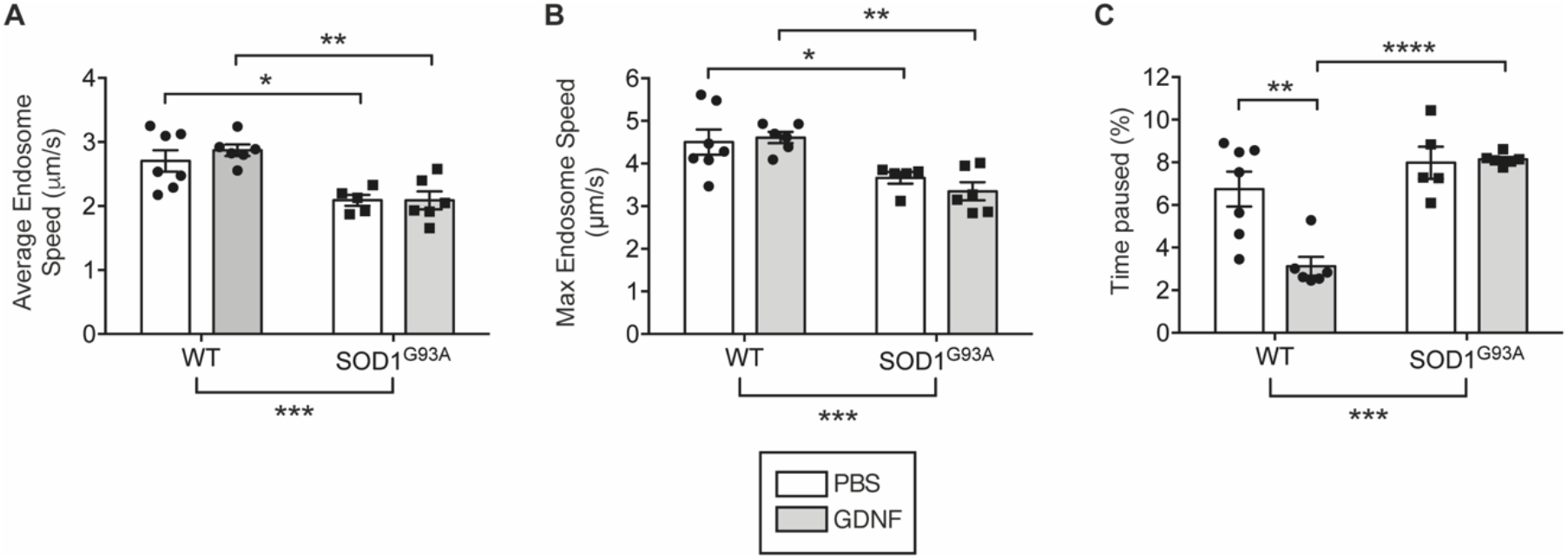
GDNF does not alter *in vivo* signalling endosome transport speeds in the WT or SOD1^G93A^ sciatic nerve. **A**. Average signalling endosome speed in the sciatic nerve of P73 WT and SOD1^G93A^ mice after intramuscular injection with 25 ng GDNF or vehicle control, PBS. GDNF had no significant effect on average endosome speeds (PBS:WT vs. GDNF:WT p = 0.61, PBS:SOD1^G93A^ vs. GDNF:SOD1^G93A^ p > 0.99). SOD1^G93A^ speeds were significantly slower than WT with both PBS and GDNF treatment (* p = 0.01 and ** p = 0.003 respectively). Statistical tests: two-way ANOVA with Holm-Šídák’s multiple comparisons. **B**. Maximum signalling endosome speed. GDNF had no significant effect on maximum endosome speed. Again, SOD1^G93A^ maximum speeds were significantly lower than WT with both PBS and GDNF treatment (* p = 0.0464 and ** p = 0.0039 respectively, two-way ANOVA with Holm-Šídák’s multiple comparisons). **C**. Signalling endosome pausing (%). GDNF significantly reduced signalling endosome pausing in the WT, but not SOD1^G93A^ sciatic nerve (** p = 0.0014). There was significantly more pausing in SOD1^G93A^ mice compared to WT with GDNF treatment (**** p < 0.0001), but no significant difference with PBS treatment (p = 0.3241). Statistical tests: two-way ANOVA with Holm-Šídák’s multiple comparisons. In all analyses, there was a significant difference between genotypes (two-way ANOVA, *** p < 0.001). N = 6-7 animals. Points represent data from each individual animal. Bars represent mean ± SEM.

